# Pollinators mediate floral microbial diversity and network under agrochemical disturbance

**DOI:** 10.1101/2020.12.05.413260

**Authors:** Na Wei, Avery L. Russell, Abigail R. Jarrett, Tia-Lynn Ashman

**Affiliations:** The Holden Arboretum, Kirtland, OH 44094, USA; Department of Biological Sciences, University of Pittsburgh, Pittsburgh, PA 15260, USA; Department of Biology, Missouri State University, Springfield, MO 65897, USA

**Keywords:** Agrochemical disturbance, bacterial and fungal communities, floral microbiome, microbial network, pollinator visitation, strawberry

## Abstract

How pollinators mediate microbiome assembly in the anthosphere is a major unresolved question of theoretical and applied importance in the face of anthropogenic disturbance. We addressed this question by linking visitation of diverse pollinator functional groups (bees, wasps, flies, butterflies, beetles, true bugs and other taxa) to the key properties of floral microbiome (microbial α- and β-diversity and microbial network) under agrochemical disturbance, using a field experiment of bactericide and fungicide treatments on cultivated strawberries that differ in flower abundance. Structural equation modeling was used to link agrochemical disturbance and flower abundance to pollinator visitation to floral microbiome properties. Our results revealed that (1) pollinator visitation influenced the α- and β-diversity and network centrality of floral microbiome, with different pollinator functional groups affecting different microbiome properties; (2) flower abundance influenced floral microbiome both directly by governing the source pool of microbes and indirectly by enhancing pollinator visitation; and (3) agrochemical disturbance affected floral microbiome primarily directly by fungicide, and less so indirectly via pollinator visitation. These findings improve the mechanistic understanding of floral microbiome assembly, and may be generalizable to many other plants that are visited by diverse insect pollinators in natural and managed ecosystems.

## Introduction

Microbiomes can profoundly influence plant fitness in natural and agricultural settings (Berendsen, Pieterse, & Bakker, 2012; Bulgarelli, Schlaeppi, Spaepen, Ver Loren van Themaat, & Schulze-Lefert, 2013; Vorholt, 2012). Relative to rhizosphere (root) and phyllosphere (leaf), the anthosphere (floral) microbiome is least well studied but has the most direct impact on plant reproductive success (Aleklett, Hart, & Shade, 2014; Rebolleda-Gómez et al., 2019; Wei & Ashman, 2018). Our understanding of what governs microbiome assembly in flowers – a highly dynamic niche – has advanced only recently (Aleklett et al., 2014; Vannette, 2020). The identified drivers include, for instance, microbial dispersal mediated by pollinators (Rebolleda-Gómez & Ashman, 2019; Vannette & Fukami, 2017), floral traits that can influence niche availability (e.g. flower size, nectar volume) (Vannette, 2020; Vannette, Hall, & Munkres, 2020) and microbial source pool (e.g. flower abundance; this study), and disturbance imposed by bactericides and fungicides especially in crop plants (Bartlewicz, Pozo, Honnay, Lievens, & Jacquemyn, 2016; Schaeffer, Vannette, Brittain, Williams, & Fukami, 2017). Yet, as these different drivers have often been studied independently, we lack a clear view as to how they act together and thus their relative importance in shaping floral microbiome.

Pollinator-mediated microbial dispersal is an important determinant of floral microbiome (Rebolleda-Gómez & Ashman, 2019; Vannette, 2020; Vannette & Fukami, 2017). Pollinators facilitate microbe colonization and transport but also impact microbial communities by consuming resident microbes (Herrera, Pozo, & Medrano, 2013; Russell, Rebolleda-Gómez, Shaible, & Ashman, 2019; Vannette, Gauthier, & Fukami, 2013). The net outcome on floral microbiome may differ among pollinator taxa (or flower visitors more generally), given their different uses of flowers and preferences for floral resources (Armbruster, 2017; Kantsa et al., 2018; Wei et al., 2020) and microbes (e.g. bacteria vs. yeasts) (Good, Gauthier, Vannette, & Fukami, 2014; Schaeffer, Rering, Maalouf, Beck, & Vannette, 2019), as well as different likelihoods of dispersing microbes based on level of interaction with flowers or flower compartments (Russell et al., 2019) or of bringing different microbes to the flowers owing to their remarkably different lifestyles that could affect associated microbes (Engel et al., 2016; Kwong et al., 2017). Research on the effect of pollinators on floral microbiome has been focused on a few pollinator taxa (e.g. honeybees, bumblebees and hummingbirds) (Good et al., 2014; Schaeffer et al., 2019; Vannette et al., 2013), while most plant species are pollinated by diverse groups of pollinator species (Kantsa et al., 2018; Wei et al., 2020). Thus significant knowledge gaps exist as to how different pollinators govern floral microbiome assembly, and further how the effect of pollinators is modulated by other factors that can influence pollinators and the floral microbiome, such as floral traits (e.g. flower abundance) and external disturbance of bactericides and fungicides.

Floral traits can influence floral microbiome directly by affecting niche availability and imposing habitat filtering for microbes (Rebolleda-Gómez & Ashman, 2019; Vannette, 2020; Wei & Ashman, 2018) and indirectly by affecting pollinator visits (Armbruster, 2017; Kantsa et al., 2018; Wei et al., 2020) that can mediate microbial dispersal. In particular, flower abundance has rarely been studied in floral microbiome but can potentially influence a given flower’s microbial community by affecting the source pool of microbes hosted by neighboring flowers. Such source pool of microbes is dynamic due to rapid habitat extinction (short flower longevity) and temporally shifted habitat availability (changes in flower abundance over time) (Aleklett et al., 2014). Thus flower abundance may directly influence the abundance and richness of microbes that colonize individual flowers independently of pollinators (e.g. by wind or rain splash). In addition, flower abundance can signal resource availability to pollinators (Benadi & Pauw, 2018; Wei et al., 2020), and thus large displays attract more visits and can potentially affect microbiome indirectly by enhancing microbial dispersal to flowers within the display. Therefore, flower abundance and pollinators act together in shaping floral microbiome, and their independent and interactive effects on floral microbiome remain to be quantified.

Theory predicts that disturbance alters niche availability, the nature of species interactions (positive, facilitation or negative, competition) and species composition of communities (Mouillot, Graham, Villeger, Mason, & Bellwood, 2013). For microbiomes in an agricultural setting, an apparent disturbance can come from bactericides and fungicides that broadly or selectively target microbial taxa (Matson, Parton, Power, & Swift, 1997; McManus, Stockwell, Sundin, & Jones, 2002). Chemically-mediated disturbances are becoming more common in the Anthropocene as agrochemical use is increasing, and can reduce the abundance and richness of plant-associated microbes, shift microbial composition (Bartlewicz et al., 2016; Schaeffer et al., 2017) and potentially rewire microbe–microbe interactions via physical and metabolic mechanisms (e.g. habitat sharing via fungal hyphae and bacterial biofilm and by-product cross-feeding) (Deveau et al., 2018; Frey-Klett et al., 2011; Goldford et al., 2018; Smith, Shorten, Altermann, Roy, & McNabb, 2019). Agrochemical disturbance has also shown off-target effects on pollinators including altering their visitation patterns (Fisher, Coleman, Hoffmann, Fritz, & Rangel, 2017; Park, Blitzer, Gibbs, Losey, & Danforth, 2015; Stejskalová, Konradyová, Suchanová, & Kazda, 2018). Thus, agrochemical disturbance of bactericides and fungicides likely affects flower microbiome both directly (by reducing targeted microbial taxa) and indirectly (by influencing pollinator visitation). It remains, nevertheless, a major unresolved question whether the effect of agrochemical disturbance is generalizable across taxonomically diverse pollinators and between bacterial and fungal constituents of floral microbiome.

Here we used a field experiment to quantify how insect pollinators govern the key properties of floral microbiome (i.e. microbial α- and β-diversity and microbial network) under agrochemical disturbance (bactericide and fungicide; Figure 1). We focused on the epiphytic floral microbiome of cultivated strawberry (*Fragaria* ×*ananassa*). Strawberry crops are commercially important worldwide (FAOSTAT, 2020) and rely heavily on pesticide use for productivity (Environmental Working Group, 2020). While the plant is capable of autonomous self-pollination, visitation by a wide array of insects is essential for pollination success and marketable fruit production (Klatt et al., 2014). Thus strawberry represents a model system to investigate how agrochemicals and pollinators affect microbiome in flowers. Rather than manipulating pollinator access (e.g. Rebolleda-Gómez & Ashman, 2019; Vannette & Fukami, 2017), we linked the natural variation in visitation of different functional groups of pollinators (bees, wasps, flies, butterflies, beetles, true bugs and other taxa) to bacterial and fungal α- and β-diversity and microbial network in flowers. To minimize habitat filtering due to host variation, our experiment used clonal replicates of four genotypes of strawberry. The four genotypes differ dramatically in flower abundance but are similar in other floral traits. To test for the effect of agrochemical disturbance, we performed a replicated factorial design using broad-spectrum bactericide and fungicide at the agricultural application rate (Figure 1a). By doing so, we first characterized the communities of visiting pollinators and floral microbiota (bacteria and fungi), and examined separately how they responded to agrochemical disturbance and plant genotype. We further constructed microbial correlation networks to infer intra-kingdom (bacteria–bacteria and fungi–fungi) and inter-kingdom (fungi–bacteria) interactions among microbes and the network centrality of floral microbiome. Lastly, to link pollinator communities to microbial communities, we used structural equation modeling to quantify how agrochemical disturbance and genotypic variation in flower abundance affected the visitation of pollinator functional groups and thereby together affected microbiome properties (α- and β-diversity and network centrality; Figure 1b). These analyses revealed that (1) pollinator visitation influenced floral microbiome with different pollinator functional groups affecting different properties of floral microbiome; (2) flower abundance strongly affected floral microbiome both directly and indirectly via enhancing pollinator visitation; (3) agrochemical disturbance affected floral microbiome primarily directly, and the effect was caused by fungicide rather than bactericide.

**Figure 1.**
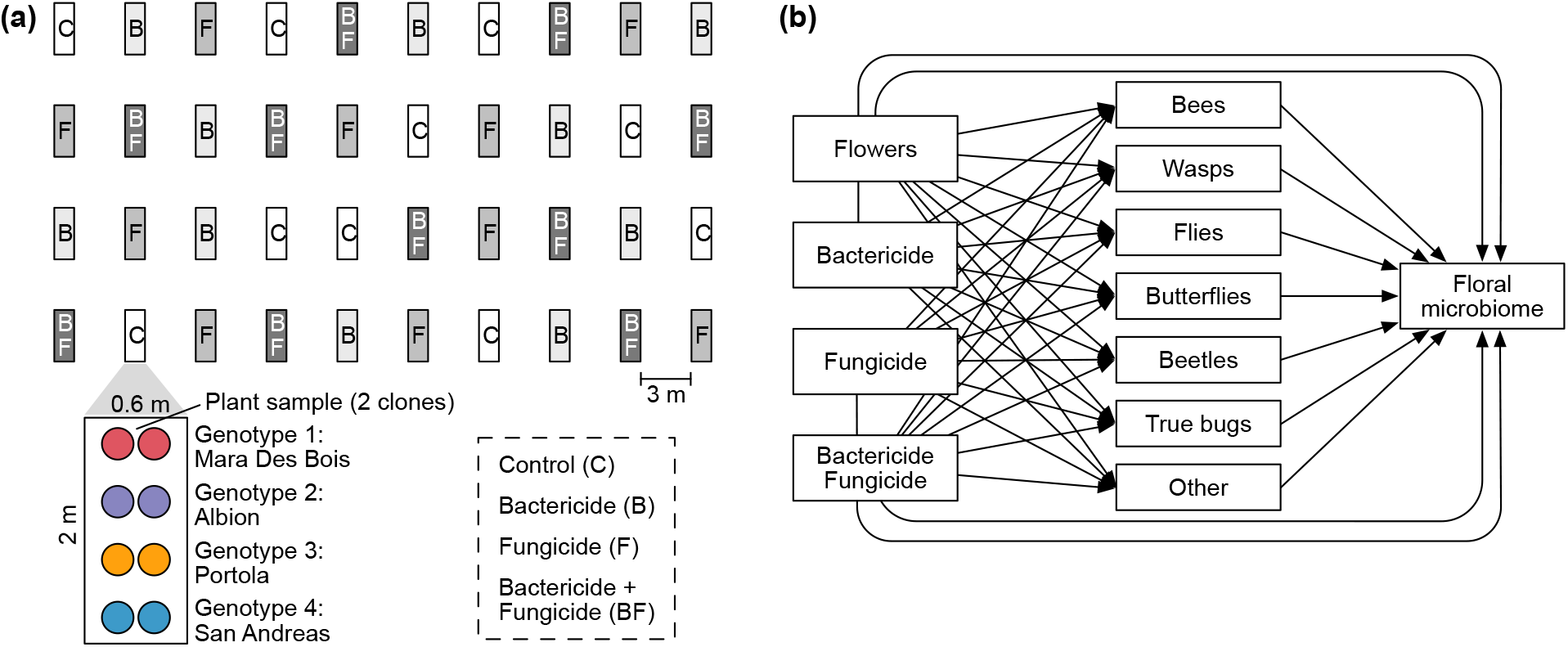
Summary of experimental design and hypotheses in structural equation modeling. (a) Completely randomized, replicated factorial design was used to examine how pollinators influence floral microbiome under agrochemical disturbance. Each treatment had 10 replicates (‘blocks’). Each block consisted of four strawberry genotypes, with two clonal plants per genotype (referred to as one plant sample) to increase floral display for pollinators. Blocks were spaced to minimize potential treatment spillover. (b) Hypothesized effects of flower abundance and agrochemical disturbance on pollinator visitation and floral microbiome. Although floral microbiome may also influence pollinator visitation, this effect was unlikely in this study (see main text) and thus one-way arrows were used to represent the effect of pollinator visitation on floral microbiome. Floral microbiome was characterized using bacterial and fungal α- and β- diversity and microbial network centrality here.

## Materials and Methods

### Experimental design

#### Strawberry plant system and factorial experimental design

We used four day-neutral strawberry cultivars (Mara Des Bois, Albion, Portola, San Andreas; Nourse Farms, Whately, MA) that show pronounced differences in flower production (Figure 2a). For each genotype (cultivar), we planted 80 clonal ’bare root’ plants in 2.4 L pots filled with Fafard 4 mix (Sun Gro Horticulture, Agawam, MA) on May 3, 2018. These plants were grown at 24°C day/18°C night temperatures 12-h days in a greenhouse at the University of Pittsburgh for 2 wk until they were transported to a field at the Pymatuning Laboratory of Ecology, Linesville PA (41.57011°N, 80.45619°W). Potted plants grew and accumulated natural microbiota for 5 wk prior to the start of a 4-wk experiment of agrochemical treatment, pollinator observation, flower abundance survey and floral microbiome collection from June 24 to July 21, 2018. The factorial design of agrochemical treatments included water control (’C’), bactericide (’B’, oxytetracycline, 150 ppm; Sigma-Aldrich, St. Louis, MO), fungicide (’F’, azoxystrobin, Heritage, 0.2 lb per acre; Syngenta, Greensboro, NC), and bactericide and fungicide (’BF’). The four treatments were applied at the ’block’ level (*N* = 40 with 10 blocks per treatment, Figure 1a). Each experimental block consisted of four pairs of clones (one pair of each genotype) and the location of pairs within block was randomized. Pairs were used to ensure floral display for pollinators and were referred to as one ’plant sample’ hereafter (Figure 1a). Treatments were applied weekly using a backpack sprayer, with plant samples receiving an equal volume across treatments each week (‘C’: 25 mL water per plant sample; ‘B’: 12.5 mL water and 12.5 mL bactericide; ‘F’: 12.5 mL water and 12.5 mL fungicide; ‘BF’: 12.5 mL bactericide and 12.5 mL fungicide). Plants were protected from large herbivores using fencing with buffer distance (10 m) to the site.

**Figure 2.**
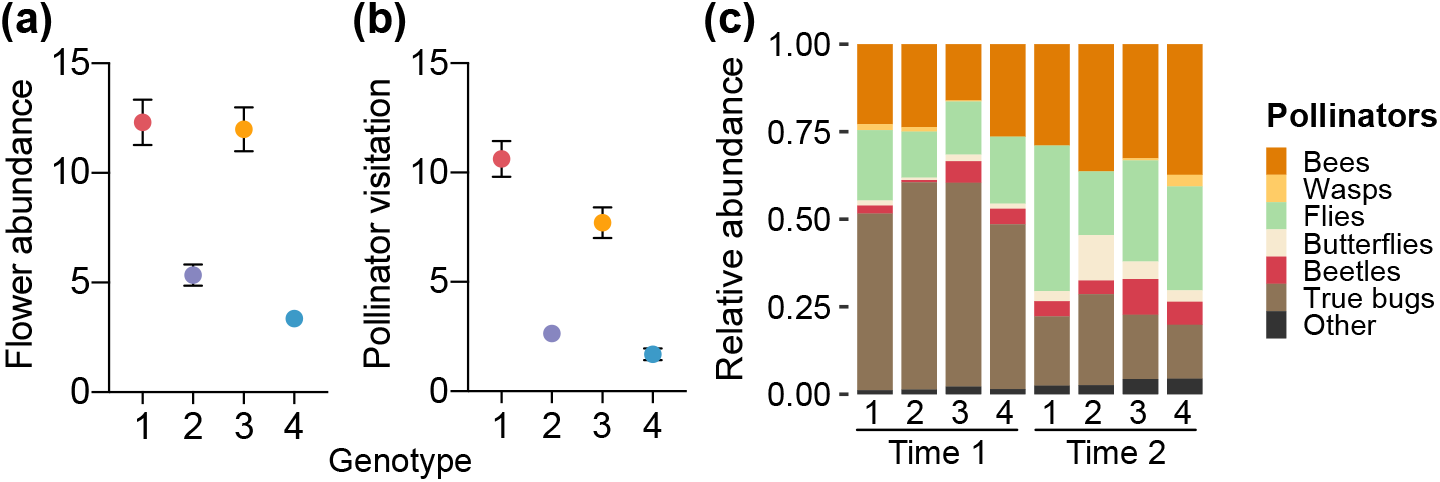
Plant genotypes differ in flower abundance and visting pollinators. Genotypic variation in (a) flower abundance matched (b) pollinator visitation (i.e. number of visiting pollinators per ∼36 min) per plant sample during a 2-wk time period. (a and b) The least-squares means are plotted with error bars (1 SE) after controlling for agrochemical treatment and time period (the first vs. last 2 wk, Time 1 vs. Time 2) in generalized linear mixed models (Tables S1 and S2). (c) Pollinator community composition changed significantly over the two time periods and among genotypes (Table S2). The four strawberry genotypes were: Mara Des Bois (1), Albion (2), Portola (3), and San Andreas (4).

During the 4-wk experiment, due to temporal dynamics in flower production and pollinator visitation between the first 2 wk and last 2 wk (Figures S1 and S2), we divided the experiment into the corresponding two time periods (Time 1 and Time 2).

#### Pollinator observation

Pollinator observations were conducted by the same observer four days each week between 0900–1700 h. The observer spent 2 min per block for all 40 blocks in a single observation round. At each block, all flowers were observed simultaneously and the number of pollinators (bees, wasps, flies, butterflies, beetles, true bugs and other taxa) were recorded for each plant sample. Each day 1–3 rounds of observations were completed with the order of block visits systematically alternated each round. As a result, we spent 2874 min total for pollinator observations in this experiment, and reported the visitation data as the number of visiting pollinators per plant sample per ∼36 min during each 2-wk time period.

#### Flower abundance survey

We scored the number of flowers including both open flowers and flower buds of each plant sample once per week, and reported flower abundance data as the sum of flowers per plant sample per 2-wk time period. We also confirmed that the four genotypes differed significantly in flower abundance (Figure 2a; SI Methods) and not other traits (e.g. flower size and pollen production, Figure S1; SI Methods) that reflect attraction and resources for pollinators (Ashman, 2000).

#### Floral microbiome collection and sequencing

Two flowers were collected from each plant sample each week two days post agrochemical application. Following Wei and Ashman (2018), flowers (without pedicel) were collected into a sterile 15 mL centrifuge tube using ethanol-rinsed forceps and were transferred to a −20 °C freezer within two hours. For microbial DNA extraction, we pooled flowers standardized by size (*N* = 1–4) per plant sample per 2-wk time period and obtained 246 total samples. Epiphytic microbes were collected by sonicating flowers in 3 mL phosphate-buffered saline at 40 kHz for 10 min and vortexing for 5 min, and were pelleted by centrifuging at 13,300 rpm for 5 min. Microbial DNA was extracted using Quick-DNA Fecal/Soil Microbe Kits (Zymo Research, Irvine, CA). Two negative controls without flower samples were included in the process of microbe isolation and DNA extraction. Samples and negative controls were sent to Argonne National Laboratory for bacterial (16S rRNA V5–V6 region, 799f–1115r primer pair) (Redford, Bowers, Knight, Linhart, & Fierer, 2010) and fungal (ITS1f–ITS2) (Smith & Peay, 2014) library preparation. Because the negative controls failed in PCRs in library preparation, the 246 samples were sequenced on two lanes of Illumina MiSeq paired-end 250 bp.

### Bioinformatic and statistical analyses

#### Microbial sequence processing

Demultiplexed paired-end (PE) reads were used for detecting bacterial and fungal amplicon sequence variants (ASVs) using package DADA2 v1.12.1 (Callahan et al., 2016) in R v3.6.0 (R Core Team, 2019) and QIIME 2 v2019.4 (Bolyen et al., 2019). For bacterial ASV analysis in DADA2, PE reads were trimmed and filtered [truncLen = c(245, 245), trimLeft = c(10, 0), maxN = 0, truncQ = 2] after initial quality inspection. Then end-specific variants were identified after taking into account sequence errors, prior to joining the PE reads (minOverlap = 20, maxMismatch = 4) for ASV detection. Bacterial ASVs were further filtered against chimeras and assigned with taxonomic identification based on the SILVA reference database (132 release) implemented in DADA2. For fungal ASV analysis in DADA2, the PE reads were first screened to remove potential primer contaminations, and then subject to the ASV detection processes from trimming and quality filtering [truncLen = c(200, 200), maxN = 0, truncQ = 2] to chimera removal, as described for bacteria. Fungal taxonomic assignment was conducted based on the UNITE reference database (v8.0 dynamic release) using QIIME 2.

Bacterial and fungal ASV tables were further filtered separately before conversion into microbial community matrices using package phyloseq (McMurdie & Holmes, 2013). First, we removed non-focal ASVs (Archaea, chloroplasts and mitochondria). Second, we filtered out low-depth samples (<100 reads; *N* = 13 and 8 samples for bacterial and fungal data set, respectively). Third, we normalized per-sample reads to the same number (i.e. the median reads, 18788 and 37516, bacterial and fungal data set, respectively) following Wei and Ashman (2018). Lastly, we removed low-frequency ASVs (<0.001% of total observations). The final community matrices consisted of 1237 and 1165 ASVs for bacteria (*N* = 223) and fungi (*N* = 240), respectively.

#### Pollinator visitation and community composition

To evaluate how pollinator visitation responded to agrochemical treatment, genotype and time period, we examined pollinator fauna as a whole and individual functional groups using zero-inflated generalized linear mixed models (zGLMMs). In each zGLMM, the predictors included treatment (levels: C, B, F and BF), genotype (1: Mara Des Bois; 2: Albion; 3: Portola; 4: San Andreas) and time period (Time 1 vs. Time 2), with random effects being block and plant sample (due to repeated measurements over time) using package glmmTMB (Brooks et al., 2017). Negative binomial errors were used in zGLMMs and model fits were confirmed using package DHARMa (Hartig, 2019). Statistical significance (type III sums of squares) and least-squares means (LS-means) of predictors were assessed using packages car (Fox & Weisberg, 2011) and emmeans (Lenth, 2019).

To evaluate how the composition of the pollinator communities responded to agrochemical treatment, genotype and time period, we converted the pollinator functional group matrix into proportional data, and conducted permutational multivariate analysis of variance (PERMANOVA; predictors: treatment, genotype and time period; random effect: block) using package vegan (Oksanen et al., 2019). Once significant predictors were detected, we further identified the specific functional groups underlying pollinator community variation caused by the predictors using random forest (RF) classification in packages caret (Kuhn, 2019) and randomForest (Liaw & Wiener, 2002). The RF classification models were run for the full data with 1000 trees, and the number of randomly selected variables (i.e. functional groups) at each split of a decision tree was optimized using 10-fold cross validation in caret. RF model performance was assessed using out-of-bag (OOB) error. The set of important variables (functional groups) were selected using backwards variable elimination with package varSelRF (Diaz-Uriarte, 2007), and their relative importance was evaluated using mean decrease in classification accuracy.

#### Floral microbial α- and β-diversity

To evaluate how microbial α-diversity responded to agrochemical treatment, genotype and time period, we calculated Shannon diversity for bacterial and fungal communities separately using vegan, and performed general linear mixed models (LMMs) using package lme4 (Bates, Machler, Bolker, & Walker, 2015). The predictors included treatment, genotype and time period as well as their two-way interactions, and the random effects included block and plant sample. The response variable was power transformed to improve normality, with the optimal power parameter determined using the Box–Cox method in package car. Statistical significance of predictors in LMMs was evaluated using package lmerTest (Kuznetsova, Brockhoff, & Christensen, 2017).

Microbial β-diversity (Bray–Curtis dissimilarity) was evaluated using PERMANOVA and constrained principal coordinates analysis (cPCoA) in vegan for bacterial and fungal communities separately. To assess the significance of the main effects, PERMANOVA and cPCoA included treatment, genotype and time period with block random effect. To assess the significance of two-way interactions, PERMANOVA and cPCoA included both the main effects and their interaction terms. Once significant predictors were identified, random forest classification was used to detect the set of important ASVs underlying bacterial or fungal community variation caused by the predictors, as described for pollinator community composition.

#### Structural equation models (SEMs) linking pollinators to microbial diversity

To evaluate how pollinators influenced microbial diversity under agrochemical disturbance, we conducted SEMs to link flower abundance and agrochemical treatment to visitation by pollinators of specific functional groups to α- and β-diversity of bacteria or fungi in the floral microbiome (Figure 1b). The SEMs included flower abundance as genotypic variation was only significant in this trait (Figure 2a) and not the others (Figure S1). The SEMs were conducted on data collected across both time periods but did not include time period as an exogenous (independent) variable, because time-period variation was defined and reflected by variation in flower abundance and pollinator visitation (Tables S1 and S2). For exogenous categorical variables of treatments, we coded the control treatment as the reference level so that the effects of all other treatments were relative to the control. The endogenous (dependent) variable of the α-diversity metric (Shannon diversity) was power transformed as in section *Floral microbial α- and β-diversity* (power parameter = 1 and 2 for bacteria and fungi, respectively). The endogenous variable of β-diversity used the first axis of the cPCoA (see section *Floral microbial α- and β- diversity*) that accounted for 25% and 60% of total variation in bacterial and fungal communities, respectively. Model estimation used robust maximum likelihood with Satorra-Bentler scaled χ^2^ that can accommodate nonnormality using package lavaan (Rosseel, 2012). Model fits were confirmed (comparative fit index, CFI > 0.9; root mean squared error of approximation, RMSEA, the lower bound of 90% confidence interval < 0.05; standardized root mean squared residual, SRMR < 0.1) (Kline, 2015). We conducted the SEMs using the original and square-root transformed pollinator visitation data separately, which yielded qualitatively similar results but the former exhibited better model fits and thus were reported here. Standardized coefficients (*r*) were indicated for paths with notable effects (*P* < 0.1).

#### Microbial networks and SEMs linking pollinators to network centrality

Microbial correlation networks were constructed to infer intra- and inter-kingdom interactions among microbes in flowers, and how pollinators influenced microbial network centrality under agrochemical disturbance. We focused on two centrality metrics (degree and eigenvector). In a network, degree centrality describes a microbe’s importance based on the total number of direct connections (or links) with other microbes. Eigenvector centrality weighs each connection by giving higher weights to connections to well-connected (i.e. high degree) microbes and considers both direct and indirect connections, and thus is a weighted sum of all connections of a microbe to other microbes (Bonacich, 2007). Therefore, degree and eigenvector centrality measure the network importance of microbes based on unweighted direct interactions and weighted network-wide interactions, respectively.

To build microbial networks, we merged the bacterial and fungal community matrices and normalized per-sample read numbers (to the median of the bacterial data set). This resulted in 2346 ASVs (1232 bacteria and 1114 fungi) across 218 samples that had data for both bacteria and fungi. To infer positive and negative microbe–microbe interactions, we used SparCC (Friedman & Alm, 2012) that is robust to spurious correlations among microbes due to microbiome compositionality (i.e. non-independence among ASVs due to relative abundances) using package SpiecEasi (Kurtz et al., 2015). To do this, we extracted the raw counts of these ASVs, and focused on the ASVs that were present in ≥10 samples. Networks were constructed and visualized for each treatment (based on correlation estimates ≥0.4 and ≤-0.4) using SparCC and package igraph (Csardi & Nepusz, 2006).

To infer how pollinators influenced microbial network (degree and eigenvector centrality) under agrochemical disturbance using SEMs (Figure 1b), we targeted ASVs that were responsive to (1) visitation of pollinator functional groups and (2) flower abundance based on maximal information coefficient (MIC ≥0.2) (Reshef et al., 2011) and (3) agrochemical treatments using random forest (RF) classification. MICs that can detect nonlinear associations were calculated using package minerva (Albanese et al., 2013). RF classification was performed to detect the ASVs that differed between pairwise comparisons of treatments for bacteria and fungi separately. This resulted in 238 ASVs (146 bacteria and 92 fungi) in the 218 samples for SEMs. To do this, we performed leave-one-out approach to measure network change caused by the removal of a sample. Specifically, we constructed the microbial network with and without a sample based on MIC (≥0.2) due to its nonlinear detectability. We then calculated node-level degree and eigenvector centrality for individual nodes (ASVs) using igraph. Network change in node-level centralities was calculated as the distance between two vectors of ASV centralities (i.e. with and without a sample) using package pdist (Wong, 2013) for degree and eigenvector centrality separately. Such network change reflected individual sample’s importance to microbial network centrality and was used in the SEMs.

## Results

### Pollinator visitation and community composition

The visitation of all pollinator fauna (χ^2^ = 4.7, df = 3, *P* = 0.20, zGLMM) and that of individual functional groups (Table S2) were similar among different agrochemical treatments, except butterflies and the ‘other’ category (Figure S2). Butterfly visitation tended to increase when both bactericide and fungicide were applied (BF vs. C; zGLMM, *t* = 2.15, *P* = 0.030, Figure S2b). The ‘other’ group of pollinators decreased under fungicide treatment relative to the control (F vs. C; *t* = −2.01, *P* = 0.045, Figure S2c). Different from agrochemical treatment, plant genotype strongly influenced visitation of all pollinators combined (χ^2^ = 178, df = 3, *P* < 0.001, Figure 2b) and that of individual functional groups (all *P* < 0.05, Table S2). In addition, pollinator visitation declined over time period (χ^2^ = 41.6, df = 1, *P* < 0.001, Figure S2a), especially for true bugs (Table S2). Similar to visitation, pollinator community composition also varied significantly over time period (PERMANOVA, *F* = 34.5, df = 1, *P* = 0.001, 12% of variation, Figure 2c) and among plant genotypes (*F* = 2.2, df = 3, *P* = 0.025, 2%, Figure 2c and Figure S2d), which were driven by changes in the relative abundance of true bugs and flies (random forest, OOB = 21.4%, Table S3; Figure 2c) and that of true bugs and beetles (OOB = 63.3%), respectively.

### Floral microbial α- and β-diversity and network

Agrochemical treatment significantly affected the α-diversity of fungal communities (LMM, *F* = 3.59, df = 3, *P* = 0.022, Table S4), after accounting for the effects of genotype (*F* = 4.09, df =3, *P* = 0.009, Figure S3) and time period (*F* = 2.04, df =1, *P* = 0.16). As expected, fungal Shannon diversity was reduced when fungicide was applied (F vs. C, χ^2^ = 7.16, df = 1, *P* = 0.007; BF vs. C, χ^2^ = 7.74, df = 1, *P* = 0.005, Figure 3c). This reduction was especially strong at Time 1 (treatment × time period, *F* = 4.20, df = 3, *P* = 0.007), whereas at Time 2 all treatments remained low in fungal Shannon diversity including the control (Figure S3b), likely due to the combined effect of agrochemical use and reduced flower abundance (over time period) that had a strong effect on fungal diversity (described below in SEMs).

**Figure 3.**
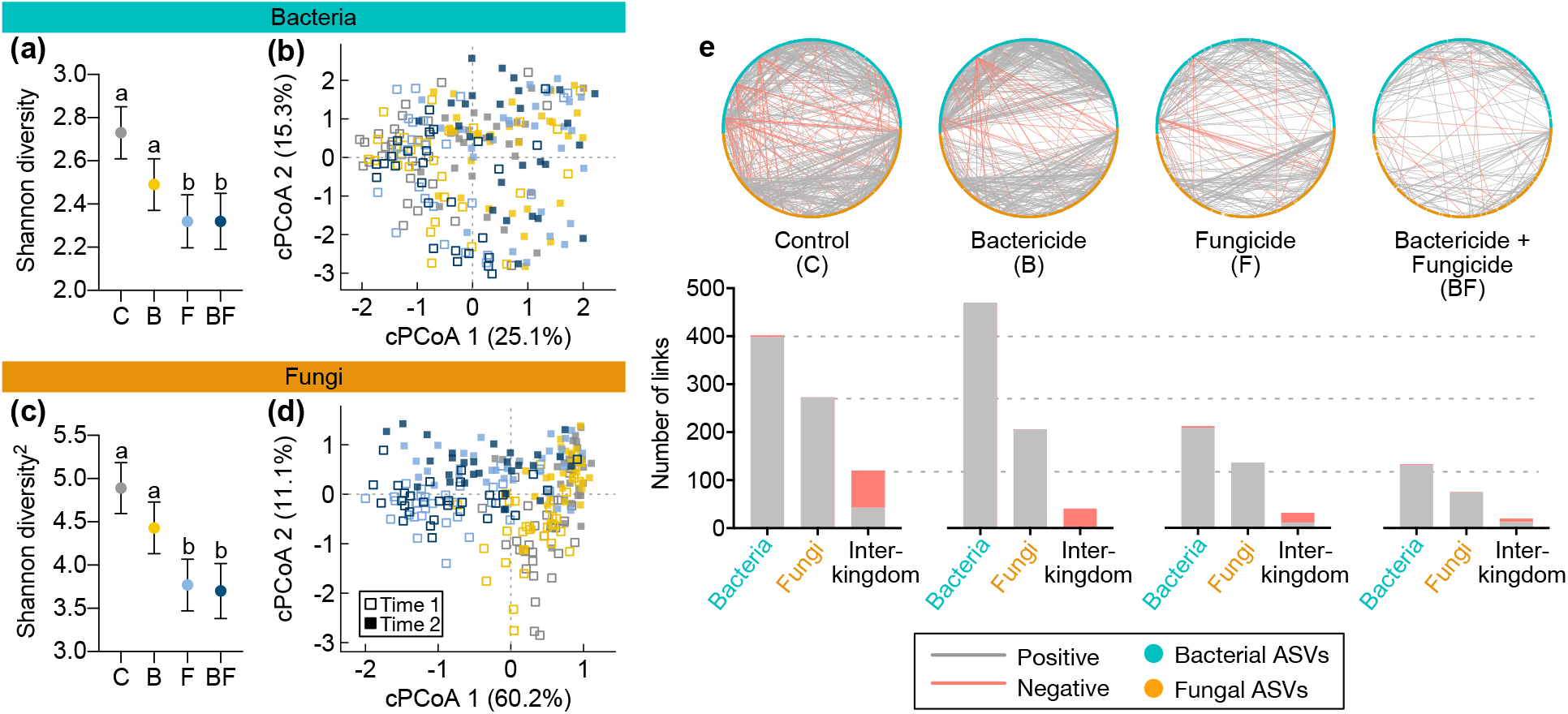
Microbial α- and β-diversity and network respond to agrochemical disturbance. The least-squares means of Shannon diversity for bacteria (a) and fungi (b; power-transformed, power parameter = 2) are plotted with error bars (1 SE) for each treatment (C, control; B, bactericide; F, fungicide; BF, bactericide and fungicide), after controlling for the effects of genotype and time period in general linear mixed models (Table S4). (b and d) Constrained principal coordinates analyses (cPCoAs) indicated microbial community separation by agrochemical treatment (color) and time period (the first vs. last 2 wk, Time 1 vs. Time 2) for bacteria (b, *N* = 223 samples) and fungi (d, *N* = 240). (e) Microbial correlation networks were based on SparCC (correlation estimates ≥0.4 or ≤−0.4). The same set of bacterial (*N* = 359) and fungal (*N* = 343) amplicon sequence variants (ASVs) are represented as nodes in the networks across treatments (e, top panel). Positive (grey) and negative (red) intra-kingdom (bacteria– bacteria and fungi–fungi) and inter-kingdom (fungi–bacteria) correlations are represented as links within the networks (e, top panel). The number of these microbial correlations (links) are summarized (e, bottom panel).

Unexpectedly, bacterial Shannon diversity was significantly reduced by fungicide (LMM, F vs. C, χ^2^ = 5.71, df = 1, *P* = 0.017; BF vs. C, χ^2^ = 5.34, df = 1, *P* = 0.021) rather than bactericide (B vs. C, χ^2^ = 1.97, df = 1, *P* = 0.16, Figure 3a and Figure S3a). The weak effect of bactericide on bacterial diversity was consistent with the observations of microbial networks (Figure 3e). Bactericide increased positive bacteria–bacteria correlations (B, 522 out of 796 total intra- and inter-kingdom links) relative to the control (399 of 795), whereas fungicide reduced positive bacteria–bacteria (232 and 146 in F and BF vs. 399 in C) and positive fungi–bacteria correlations (12 and 14 in F and BF vs. 43 in C), as well as the total links (432 and 254 in F and BF vs. 795 in C).

Similar to α-diversity, fungal community composition was affected by agrochemical treatment, which explained the largest source of variation in fungal communities (PERMANOVA, 15%, *F* = 15.2, df = 3, *P* = 0.001, Table S4) relative to time period (5%, *F* = 15.9, df = 1, *P* = 0.001) and plant genotype (3%, *F* = 2.7, df = 3, *P* = 0.001). The cPCoA further revealed that fungicide caused the divergence of fungal communities of F and BF treatments from the control (along the first axis, cPCoA1, Figure 3d). This divergence was driven by a small set of ASVs (F vs. C, *N* = 40, OOB = 11.5%; BF vs. C, *N* = 27, OOB = 9.2%), among which Basidiomycota and Ascomycota fungi exhibited the largest importance (Figure S4 and Table S5). In particular, *Rhodotorula graminis* (yeast) was significantly enriched, and *Sporobolomyces phaffii* (yeast) and *Cladosporium delicatulum* (mold) were significantly depleted in F and BF treatments relative to the control (adjusted *P* for multiple comparisons < 0.001, Table S5).

Different from fungi, bacterial communities were primarily segregated over time period along cPCoA1 (*F* = 6.96, df = 1, *P* = 0.001, Figure 3b; PERMANOVA, *F* = 7.62, *P* = 0.001, Table S4). Albeit subtler than fungi (Figure 3b,d), bacterial communities varied among agrochemical treatments (cPCoA, *F* = 1.38, df = 3, *P* = 0.017; PERMANOVA, *F* = 1.44, *P* = 0.015, Table S4), with all other treatments (B, F and BF) significantly deviated from the control (PERMANOVA, pairwise comparisons, all *P* < 0.05; data not shown). Such deviation was driven by a small set of ASVs (*N* = 8, 14 and 3 for B, F and BF, respectively, Table S5), among which two *Hymenobacter* ASVs were most influential (Figure S5) and significantly depleted in other treatments relative to the control (adjusted *P* < 0.001, Table S5).

### Pollinators influenced floral microbiome

SEMs revealed that pollinator functional groups varied in their influence on different properties of floral microbiome (α- and β-diversity and network centrality), independent of the significant direct effects of flower abundance and agrochemical treatments (Figure 4). Bee visitation had a notable positive effect on bacterial Shannon diversity (*r* = 0.12, *P* = 0.070, Figure 4a) but a negative effect on fungal Shannon diversity (*r* = −0.13, *P* = 0.068, Figure 4b), whereas butterfly visitation tended to reduce bacterial Shannon diversity (*r* = −0.08, *P* = 0.065). In contrast to α-diversity, the β-diversity of microbial communities (cPCoA1) was influenced by a different set of pollinators (Figure 4c,d). Visitation of flies and butterflies showed contrasting effects relative to true bugs on bacterial (*r* = 0.15 and 0.11 vs. −0.17, *P* = 0.015, 0.046 and 0.036, respectively, Table S6) and fungal community composition (*r* = 0.19 and 0.12 vs. −0.20, *P* < 0.001, *P* = 0.011 and *P* < 0.001, respectively). Different from α- and β-diversity, pollinator visitation did not affect microbial network degree centrality (Figure 4e). But visitation of bees and flies influenced eigenvector centrality (*r* = −0.09 and −0.17, *P* = 0.083 and 0.014, respectively, Figure 4f).

**Figure 4.**
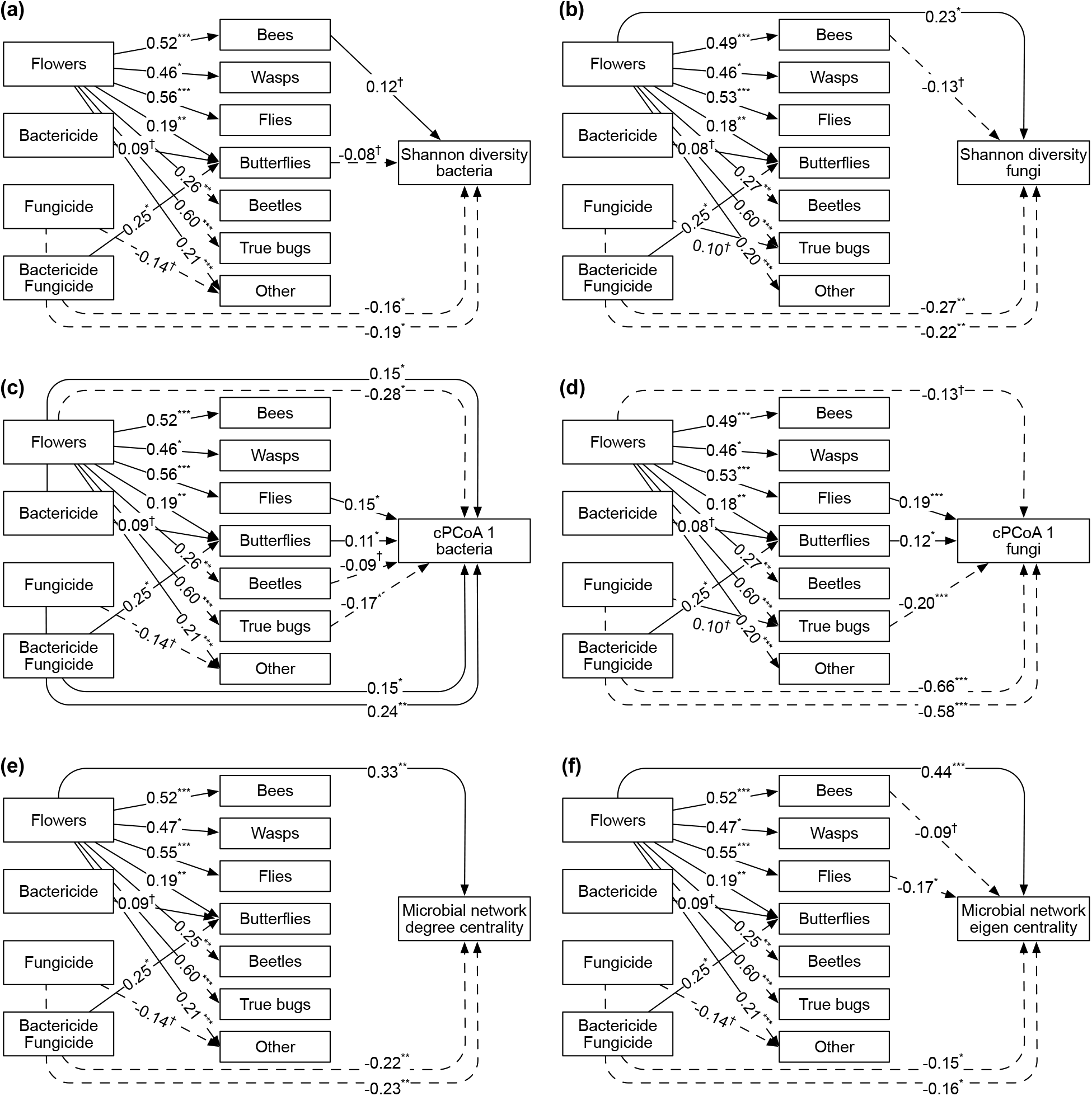
Structural equation models of flower abundance and agrochemical disturbance explaining pollinator visitation and floral microbiome. Arrows indicate notable positive (solid) and negative (dashed) relationships: †*P* < 0.10; \**P* < 0.05; \*\**P* < 0.01; \*\*\**P* < 0.001. Numbers adjacent to arrows indicate standardized path coefficients. Agrochemical treatments were coded using the control treatment as the reference level in the structural equation models (SEMs). The α- and β-diversity of bacterial and fungal communities used Shannon diversity (a and b) and the first axis of constrained principal coordinates analysis (cPCoA1, c and d; see also Figure 3), respectively. Microbial network was built on a reduced set of amplicon sequence variants (ASVs; *N* = 238, 146 bacteria and 92 fungi) using maximal information coefficient (MIC ≥ 0.2; see Methods for details). (e) The degree centrality and (f) eigenvector centrality measure network importance of individual ASVs based on the number of direct interactions and the weighted sum of both direct and indirect interactions in a network, respectively. Details in SEM model fit and parameter estimation are described in Table S6.

### Flower abundance affected floral microbiome directly and indirectly

SEMs revealed that flower abundance affected floral microbiome both directly and indirectly via influencing pollinator visitation (Figure 4). Flower abundance had a direct positive effect on fungal Shannon diversity (*r* = 0.23, *P* = 0.024, Figure 4b), and also directly affected microbial community composition (bacteria, *r* = −0.28, *P* = 0.011, Figure 4c; fungi, *r* = −0.13, *P* = 0.088, Figure 4d) and network centrality (degree, *r* = 0.33, *P* = 0.006, Figure 4e; eigenvector, *r* = 0.44, *P* < 0.001, Figure 4f). In addition, flower abundance affected microbiome indirectly by increasing pollinator visitation consistently across all functional groups (all *P* < 0.05, Figure 4). In particular, bacterial Shannon diversity was affected by flower abundance only indirectly via pollinators [bees, *r* = 0.06 (0.52 × 0.12); butterflies, *r* = −0.02 (0.19 × −0.08), Figure 4a], whereas fungal Shannon diversity was affected by flower abundance both directly and indirectly [bees, *r* = −0.06 (0.49 × −0.13), Figure 4b], similar to other properties of microbiome (community composition and eigenvector centrality). The indirect effect of flower abundance was similar between bacterial community composition [flies, *r* = 0.08 (0.56 × 0.15); butterflies, *r* = 0.02 (0.19 × 0.11); beetles, *r* = −0.02 (0.26 × −0.09); true bugs, *r* = −0.10 (0.60 × −0.17), Figure 4c] and fungal community composition [flies, *r* = 0.10 (0.53 × 0.19); butterflies, *r* = 0.02 (0.18 × 0.12); true bugs, *r* = −0.12 (0.60 × −0.20), Figure 4d]. For network eigenvector centrality (Figure 4f), the indirect effect of flower abundance was via bees [*r* = −0.05 (0.52 × −0.09)] and flies [*r* = −0.09 (0.55 × −0.17)].

### Agrochemical disturbance affected floral microbiome primarily directly by fungicide

While the significant effect of agrochemical treatment on microbiome described above (LMMs, PERMANOVA/cPCoA and network links, Figure 3) reflected the overall effect of agrochemical treatment, SEMs dissected the overall effect into direct vs. indirect effect (Figure 4). The indirect effect of agrochemical treatment (B, F and BF) via pollinators was weak, because pollinator visitation was rarely affected by agrochemical treatment except for butterflies (*r* = 0.25, *P* < 0.05, Figure 4) and the ‘other’ group (*r* = −0.14, *P* < 0.10) consistent with the zGLMMs results above, as well as true bugs (SEMs, *r* = 0.10, *P* < 0.10; zGLMM, *t* = 1.67, *P* = 0.097). The indirect effect of agrochemical treatment was only present via butterflies on bacterial Shannon diversity [B, *r* = −0.01 (0.09 × −0.08); BF, *r* = −0.02 (0.25 × −0.08), Figure 4a] and community composition [B, *r* = 0.01 (0.09 × 0.11); BF, *r* = 0.03 (0.25 × 0.11), Figure 4c], and on fungal community composition via butterflies [B, *r* = 0.01 (0.08 × 0.12); BF, *r* = 0.03 (0.25 × 0.12), Figure 4d] and true bugs [F, *r* = −0.02 (0.10 × −0.20)]. In contrast to the weak indirect effect, the direct effect of agrochemical treatment was strong for all measured microbiome properties and for both bacteria and fungi (all *P* < 0.05, Figure 4). Similar to the overall effect revealed by other methods (LMMs, PERMANOVA/cPCoA and network links, Figure 3), the direct effect of agrochemical treatment on microbiome was driven by fungicide (F and BF, Figure 4).

## Discussion

Disentangling potentially interacting mechanisms in shaping microbiomes is critical for theoretical and applied advances in the principles of microbiome assembly (Hawkes & Connor, 2017). Our work demonstrated the role of taxonomically diverse insect pollinators, and their interactions with other drivers in shaping the properties of floral microbiome. Pollinator functional groups showed distinct effect on microbial α- and β-diversity and network centrality. The effect of pollinator visitation was strongly influenced by flower abundance but less so by agrochemical disturbance. Instead, agrochemical disturbance primarily from fungicide directly affected both bacterial and fungal communities. By linking these mechanisms previously studied in isolation (see refs in Introduction), our results provide an integrated understanding of the drivers and their respective importance in governing microbiome assembly in the anthosphere in the face of agrochemical disturbance.

Our results revealed that pollinator functional groups influenced bacterial and fungal communities, even after accounting for the strong direct effects of flower abundance and agrochemical disturbance. In particular, the most abundant functional groups that visited strawberry flowers (bees, flies and true bugs, Figure 2c) played the most important roles in governing microbial diversity relative to other pollinators. These abundant pollinators differed in respective effects on microbial α-vs. β-diversity (Figure 4), with bee visitation influencing α-diversity and true bugs and flies influencing β-diversity. These distinct effects may reflect different rates of microbial dispersal mediated by different functional groups. Microbial dispersal via bees, the most important insect pollinators of cultivated strawberry (Klatt et al., 2014), can be potentially extensive and thus increase α-diversity (e.g. in bacterial communities), but high rate of microbial dispersal can also homogenize microbial communities thereby contributing little to community differentiation (along cPCoA1) relative to other functional groups (Figure 4c,d). In contrast, the effect of visitation by true bugs and flies on microbial community differentiation may reflect microbial dispersal limitation mediated by these abundant functional groups. Whether dispersal limitation is caused by shorter travel distance among flowers or fewer microbes being transported and delivered by true bugs and flies relative to bees needs further investigation. In addition to the distinct effects among pollinator functional groups, the same functional group can affect bacterial and fungal communities differently. For instance, bees tended to increase bacterial but decrease fungal α-diversity. While not affecting pollinator visitation (see discussion below), resident microbes may affect whether and how much pollinators sample and consume nectar with fungi given their preference for fungi (e.g. yeasts) over bacteria (Good et al., 2014; Vannette et al., 2013). This may lead to the negative net outcome of pollinator visitation on fungal diversity but positive effect on bacterial diversity. Taken together, it suggests that diverse insect pollinators can influence the key properties of bacterial and fungal communities.

Pollinator-mediated dispersal did not act alone but interacted with other processes in driving floral microbiome. Contrary to the expectation of off-target effects on pollinators of agrochemical use (Park et al., 2015; Stejskalová et al., 2018), visitation of most functional groups did not respond to bactericide and/or fungicide. Rather pollinator visitation was strongly influenced by flower abundance that signals resource availability, in agreement with the observations across a broad range of plant lineages in natural ecosystems (Benadi & Pauw, 2018; Wei et al., 2020). Here, flower abundance influenced not only pollinator-mediated microbial dispersal but it also had direct effect on microbiome properties suggesting it influenced the source pool of microbes. The direct and indirect effect of flower abundance underscored the importance of considering this driver in the principles of microbiome assembly in flowers.

Although floral microbes have been implicated in mediating plant–pollinator interactions (reviewed in Vannette, 2020), our results showed little evidence. This is because while microbes were significantly reduced, pollinator visitation remained unaffected under fungicide disturbance. In other words, flowers with significantly altered microbiota in the field did not seem less or more attractive to pollinators. The discrepancy with previous studies likely has three reasons. First, studies to date (reviewed in Vannette, 2020) often used artificial flowers and/or nectar to ensure that changes in cues for pollinators were induced by microbes. But in natural environments many other factors including flower abundance (Armbruster, 2017; Kantsa et al., 2018; Wei et al., 2020) can also cue the presence or quality of floral resources and thereby influence pollinator visitation, which are perhaps more salient than microbes in mediating plant– pollinator interactions in nature. Second, effects on only a few pollinator taxa have been tested to date (Good et al., 2014; Schaeffer et al., 2019; Vannette et al., 2013), so it remains an open question whether and how taxonomically diverse pollinators respond to floral microbes. Lastly, previous studies that focused on one or a few microbes (reviewed in Vannette, 2020) neglected complex microbe–microbe interactions in natural microbiomes as seen here. Thus it merits further investigation whether findings based on a few microbes are generalizable to natural microbiomes consisting of thousands or more taxa of bacteria and fungi in many flowering plants.

Agrochemical disturbance from fungicide directly affected floral microbiome, consistent with the observations in other systems and/or plant organs (Bartlewicz et al., 2016; Debode, Van Hemelrijck, Creemers, & Maes, 2013; Gu et al., 2010; Schaeffer et al., 2017). Although oxytetracycline is among the most widely used bactericides (McManus et al., 2002; Vidaver, 2002), it caused non-significant reduction in bacterial α-diversity and mild changes in composition such as *Hymenobacter*, the common bacteria in strawberry flowers (Wei & Ashman, 2018). Yet, we could not rule out that it may reduce the absolute abundance of bacteria, given that sequencing-based characterization of microbiomes considered only relative abundances. In contrast to the bactericide, azoxystrobin, one of the most popular fungicides that target eukaryotes (Battaglin, Sandstrom, Kuivila, Kolpin, & Meyer, 2010), affected both fungal and bacterial communities. It is perhaps not surprising that fungicide affected fungal communities (Bartlewicz et al., 2016; Debode et al., 2013; Gu et al., 2010; Schaeffer et al., 2017). But the strong effect of fungicide on bacterial communities underscored important inter-kingdom interactions. Fungi–bacteria interactions have been found predominantly negative in rhizosphere (Duran et al., 2018), but these interactions can be context dependent and shift in directions from negative to more positive in resource-limited environments (Velez et al., 2018), such as anthosphere here (Figure 3e). The observed positive fungi–bacteria interactions likely had physical (e.g. habitat sharing via fungal hyphae and bacterial biofilm) and/or metabolic reasons (e.g. by-product cross-feeding) (Deveau et al., 2018; Frey-Klett et al., 2011). More importantly, microbe–microbe interactions were beyond pairwise but formed a complex network, leading to cascading effects of agrochemical disturbance via network links. Therefore, reducing fungi affected not only positive fungi–bacteria but also positive bacteria–bacteria interactions (Figure 3e). Overall, our results indicated complex species interactions underlying how floral microbiome responded to agrochemical disturbance.

In conclusion, the functional links between pollinator-mediated dispersal, flower abundance and agrochemical disturbance improve the mechanistic understanding of microbiome assembly in flowers. These findings may be generalizable to many other plants in natural and managed ecosystems, as we considered not only a plant species visited by diverse insect pollinators but also the common agrochemical disturbance (Battaglin et al., 2010; Vidaver, 2002) that plants may encounter. In light of agricultural intensification (Matson et al., 1997; McManus et al., 2002) and pollinator loss worldwide (Potts et al., 2010), it becomes more urgent than ever to understand how these anthropogenic changes in plant–pollinator interactions and disturbance alter microbiomes of on- and off-target plant species.

## Acknowledgements

This work was funded by the Dr. Frank J. Schwartz Early Career Research Fellowship and G. Murray McKinley Research Fund from the Pymatuning Laboratory of Ecology (NW and ALR) and UPitt Dietrich School of Arts and Sciences (TLA). We thank L. Follweiler for greenhouse and agrochemical application assistance, N. Streher, E. O’Neill, R.A. Hayes, D. Chang, N. Russo, J. Xia, E. James and N. Cullen for assistance in the field and DNA extraction, C. Richards-Zawacki, C.J. Davis and J.A. Barabas for logistic support at the PLE field station.

