## Supplemental Information for "Pollinators mediate floral microbial diversity and network under agrochemical disturbance"

**This PDF file includes:**

Methods
Figures S1 to S5

**Other Supplementary Materials for this manuscript include the following:**

Table S1 Plant genotypes differ in flower abundance but not other traits. (see separate excel file)

Table S2 The effects of individual predictors on pollinator visitation and community composition. (see separate excel file)

Table S3 Random forest classification identified specific pollinator functional groups that underlie pollinator community variation over time period and among genotypes. (see separate excel file)

Table S4 The effects of individual predictors on bacterial and fungal α- and β-diversity. (see separate excel file)

Table S5 Random forest classification identified specific ASVs that underlie microbial community variation among agrochemical treatments for bacteria and fungi separately. (see separate excel file)

Table S6 Structural equation models of microbial α- and β-diversity and network centrality. (see separate excel file)

### Methods

*Flower abundance and other traits*

In addition to flower abundance, we also measured other traits (flower size and pollen production) that reflect attraction and resources for pollinators (Ashman, 2000) at the end of the 4-wk experiment. Flower size (i.e. corolla area) was averaged over two flowers on separate inflorescences per plant sample. We counted the number of anthers per flower in a random 54 of the 160 plant samples, and collected two anthers in a 1.5 mL microcentrifuge tube for pollen counting. The anthers each tube were acetolyzed (Dafni, 1992) to a achieve a volume of 100 µL. We enumerated pollen of a 5 µL aliquot using a hemocytometer and calculated total pollen grains per flower (pollen per anther × anthers per flower).

To confirm that the genotypes differ in flower abundance but not other traits (flower size and pollen grains per flower), we used general linear models (LMs) and a generalized linear mixed model (GLMM) in R v3.6.0 (R Core Team, 2019). In the GLMM of flower abundance using package lme4 (Bates, Machler, Bolker, & Walker, 2015), the response variable was the sum of flowers per plant sample per 2-wk time period. The predictors included treatment (C, B, F and BF), time period (the first 2 wk vs. last 2 wk, Time 1 vs. Time 2) and genotype, along with the random effects (i.e. block and plant sample due to repeated measurements over time). Poisson errors were used in the GLMM and model fit was checked using package DHARMa (Hartig, 2019). For the other traits that were measured at the end of the experiment, predictors only included treatment and genotype in the LM for flower size, and included treatment, genotype, and flower size (as a covariate) in the LM for pollen grains per flower. We did not include block random effect due to the influence of missing data on model convergence, especially for traits that were measured in a subset of plant samples. Response variables of LMs were power transformed to improve normality, with the optimal power parameter determined using the Box–Cox method in package car (Fox & Weisberg, 2011). For all models, statistical significance of predictors was evaluated by type III sums of squares using package lmerTest (Kuznetsova, Brockhoff, & Christensen, 2017) for the GLMM and package car for the LMs. The least-squares means (LS-means) of predictors were estimated using package emmeans (Lenth, 2019).

These analyses confirmed that the four genotypes differed significantly in flower abundance (Figure 2a) but were similar in flower size and pollen grains per flower (Figure S1). Flower abundance decreased significantly over the two time periods (Figure S1a).

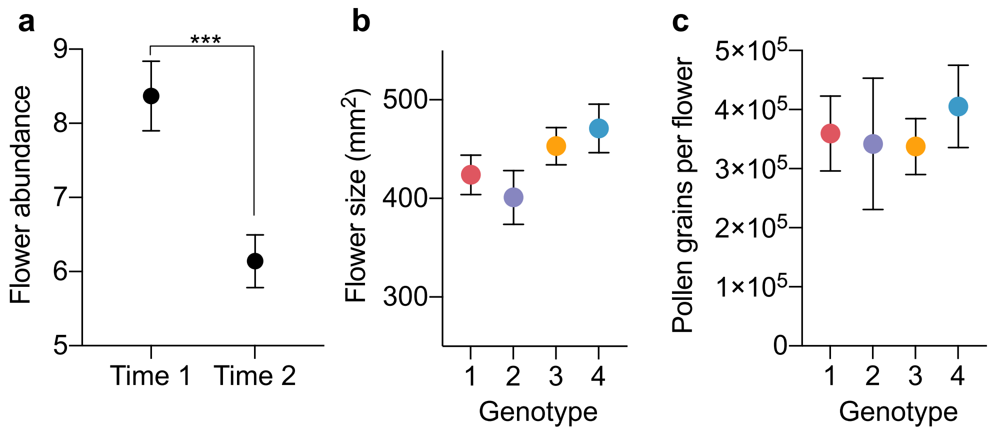


**Figure S1** Flower abundance and other traits. (a) The least-squares means (LS-means) are plotted with error bars (1 SE) for flower abundance per plant sample per 2-wk time period in the generalized linear mixed model, after controlling for the effects of genotype (1, Mara Des Bois; 2, Albion; 3, Portola; 4, San Andreas) and agrochemical treatment. The two time periods were the first 2 wk and last 2 wk (Time 1 vs. Time 2) in this study. (b) Strawberry genotypes did not differ significantly in flower size (LS-means and 1 SE), as revealed by the general linear model after controlling for agrochemical treatment. (c) Strawberry genotypes did not differ significantly in pollen production per flower (LS-means and 1 SE), as revealed by the the general linear model after controlling for flower size and agrochemical treatment. Due to the log transformation of pollen grains per flower, the LS-means were back transformed. All model details are described in Table S1.

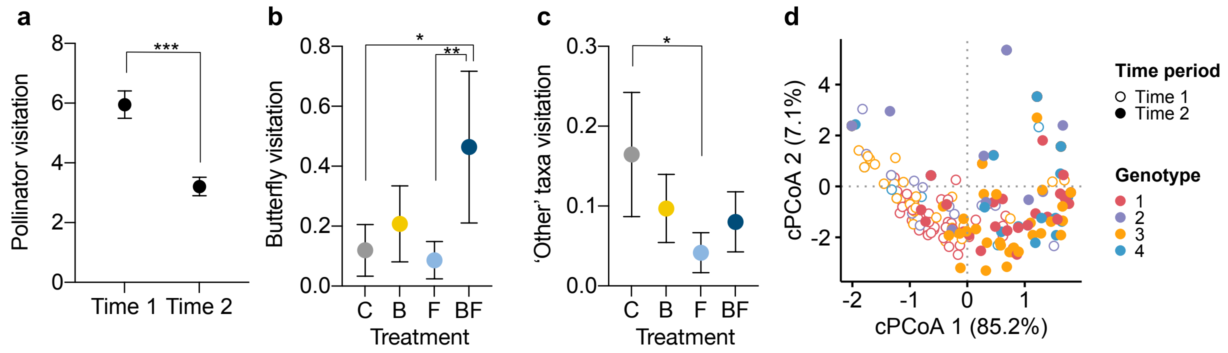


**Figure S2** Pollinator visitation and community structure. (a) The least-squares means (LS-means) are plotted with error bars (1 SE) for pollinator visiation (i.e. number of visiting pollinators per plant sample per ~36 min during a 2-wk time period), after controlling for agrochemical treatment and plant genotype in a zero-inflated generalized linear mixed model (zGLMM). (b) Buttefly visitation and (c) the 'other' taxa visitation (LS-means) differed among certain treatments (C, control; B, bactericide; F, fungicide; BF, bactericide and fungicide), after controlling for time period and plant genotype in zGLMMs. (d) The constrained principal coordinates analysis (cPCoA) showed strong pollinator community separation by time period (Time 1 vs. Time 2, the first 2 wk vs. last 2 wk) and weaker yet significant separation by plant genotype (1, Mara Des Bois; 2, Albion; 3, Portola; 4, San Andreas). In particular, genotype 4 was mostly distriubted along the positive axis of cPCoA1 and the other genotypes were more widespread. Model details are described in Table S2.

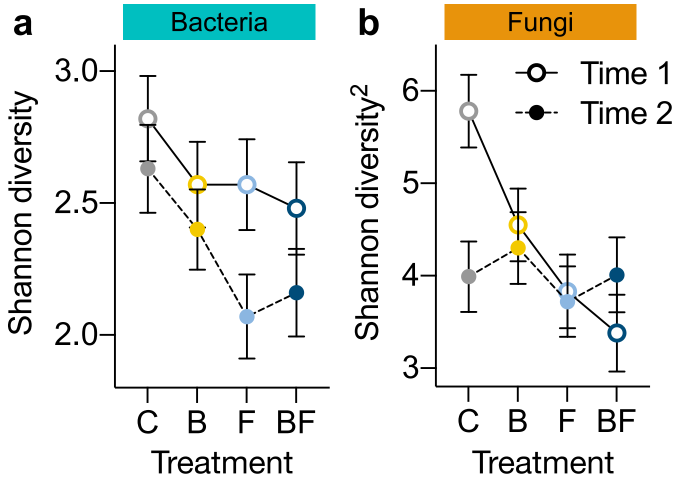


**Figure S3** Bacterial and fungal Shannon diversity responded differently to agrochemical treatment and time period. The least-squares means are plotted with error bars (1 SE) in the general linear mixed models (LMMs) for bacteria (a) and fungi (b; power transformation with power parameter = 2), after controlling for all other factors (Table S4). (a) Bacterial Shannon diversity was not significantly affected by bactericide treatment (B) at both the first 2 wk (Time 1) and last 2 wk (Time 2), but was significantly lowered by fungicide (F) and bactericide and fungicide (BF) treatments relative to the control (C) at Time 2. (b) Fungal Shannon diversity was significantly affected by agrochemical treatment and there was a significant treatment × time period interaction. Model detailed are described in Table S4.

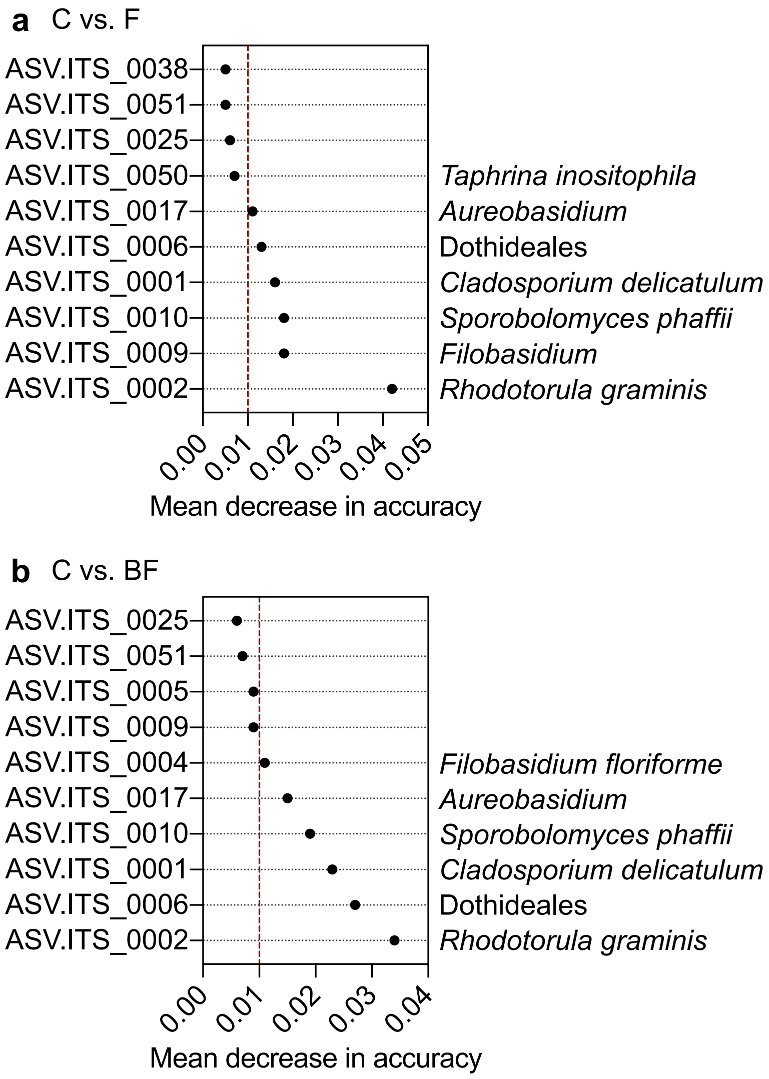


**Figure S4** Amplicon sequence variants (ASVs) underlying fungal community difference between fungicide use and the control. The selected set of important ASVs were ranked by their mean decrease in model accuracy in random forest classification. Only top ASVs (up to 10) are plotted (see the full set in Table S5). The vertical red lines indicate an arbitrary cutoff of 0.01. C, control treatment; F, fungicide treatment; BF, bactericide and fungicide treatment.

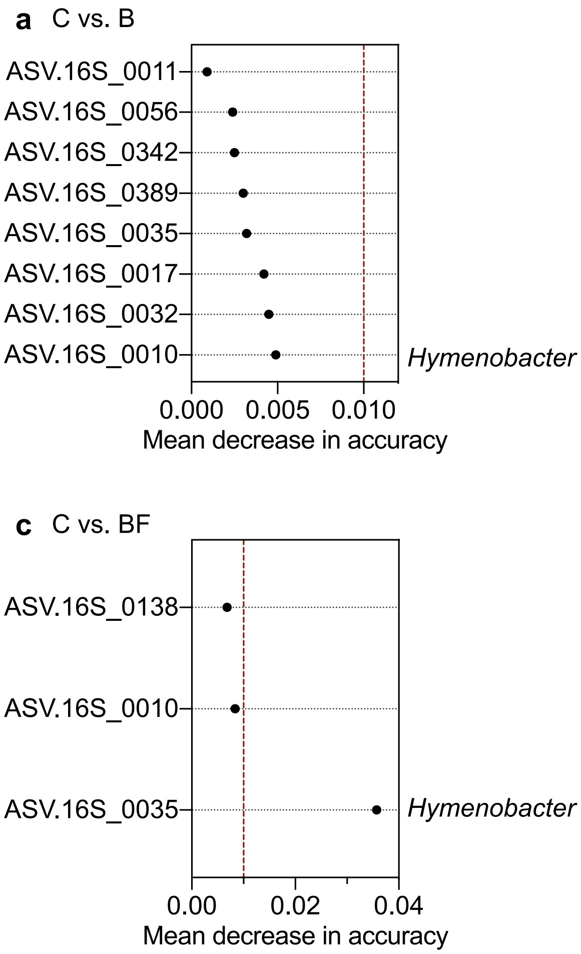


**Figure S5** Amplicon sequence variants (ASVs) underlying bacterial community difference between agrochemical use and the control. The selected set of important ASVs were ranked by their mean decrease in model accuracy in random forest classification. Only top ASVs (up to 10) are plotted (see the full set in Table S5). The vertical red lines indicate an arbitrary cutoff of 0.01. C, control treatment; B, bactericide treatment; F, fungicide treatment; BF, bactericide and fungicide treatment.
